# Detection of *Leishmania* DNA in saliva among patients with HIV/AIDS in Trang Province, southern Thailand

**DOI:** 10.1101/299297

**Authors:** Netranapha Pandey, Suradej Siripattanapipong, Saovanee Leelayoova, Jipada Manomat, Mathirut Mungthin, Peerapan Tan-ariya, Lertwut Bualert, Tawee Naaglor, Padet Siriyasatien, Atchara Phumee, Phunlerd Piyaraj

**Author notes:** Corresponding author: Phunlerd Piyaraj, MD. PhD. Department of Parasitology, Phramongkutklao College of Medicine317/5 Ratchavithi Rd, Ratchathewi, Payathai, Bangkok, 10400, Thailand.

## Abstract

Leishmaniasis is a neglected tropical disease causing opportunistic infection among patients with HIV/AIDS. The fatal form of this disease is visceral leishmaniasis (VL). DNA of *Leishmania* can be detected in saliva, for which the collection is noninvasive and requires little expertise. This study aimed to evaluate the sensitivity and specificity of a nested-PCR to amplify the Internal Transcribed Spacer 1 (ITS1) to detect *Leishmania* DNA in paired saliva and buffy coat samples of 305 Thai patients with HIV/AIDS in Trang Hospital, Trang Province, southern Thailand. For asymptomatic *Leishmania* infection among Thai patients with HIV/AIDS, the sensitivity and specificity of the nested-PCR-ITS1 in buffy coat were 73.9 and 100%, respectively. However, the sensitivity in saliva was 26.1% and specificity was 100%. Using the nested-PCR-ITS1, saliva and buffy coat samples showed positive agreement in only 52.0% of patients. Saliva tested results with the nested-PCR-ITS1 showed positive agreement with the Direct Agglutination Test (DAT) in 46.5% of patients. Only 12.1% of the samples showed positive agreement for *Leishmania* infection among all the three tests: saliva, buffy coat and DAT results. Using nucleotide sequencing, at least three species of *Leishmania* infection were identified in saliva, i.e., *L. siamensis* (n=28), *L. martiniquensis* (n=9), and *L. donovani complex* (n=1). As a result, buffy coat still appears to be a better specimen to diagnose asymptomatic VL infection among individuals with HIV. However, the use of both buffy coat and saliva together as clinical specimens would increase the sensitivity of *Leishmania* detection.

## INTRODUCTION

Leishmaniasis, a vector-borne disease, is transmitted by female phlebotomine sandflies. This neglected tropical disease (NTD) is caused by hemoflagellate intracellular parasites of the genus, *Leishmania*. (1, 2) Regarding a report by the World Health Organization, 2017, leishmaniasis is the third most important vector borne disease and second leading cause of death after malaria, (3) estimated at 700,000 to 1 million new cases and 20,000 to 30,000 deaths occur annually. (1) Leishmaniasis is endemic in 97 countries of Asia, Africa and Latin America putting 5 continents with 350 million people at risk (1)

The clinical presentation of this disease depends on the interplay between the infected *Leishmania* species and host immune responses. Hence, the clinical manifestation can be divided in three major forms: cutaneous leishmaniasis (CL), mucocutaneous leishmaniasis (ML) and visceral leishmaniasis (VL). Of these three, VL is the most severe form of this disease entailing a mortality rate of almost 100% when left untreated. VL is mainly caused by two *Leishmania* species, i.e., *L. donovani* and *L. infantum*, which are endemic in many regions of the world. Up to now, VL has been reported in 35 countries and approximately 90% of the prevalence has been reported in five countries, i.e., India, Bangladesh, Nepal, Brazil and Sudan.(4)

In Thailand, VL cases have been sporadically reported both in immunocompetent and immunocompromized patients since the first autochthonous VL case was reported in 1999.

Molecular analyses have revealed that most VL cases reported in Thailand were caused by two novel *Leishmania* species, that is, *L. martiniquensis* (MON-229, WHO code: MHOM/TH/2011/PG and MHOM/MQ/92/MAR1) and *L. siamensis* (MON-324, WHO code: MHOM/TH/2010/TR).(5) These reported cases were predominantly from southern and northern Thailand and about 40% involved patients with HIV/AIDS who constituted a high risk group for *Leishmania* infection. (5) In the last two decades, leishmaniasis has been considered an opportunistic disease among patients with HIV. Leishmaniasis has also been reported among patients with other immunosuppressed states, such as transplantation, hematological malignancies, rheumatological/connective tissue diseases and chronic inflammatory conditions. (6) Leishmaniasis and HIV co-infection is one major complications and countries facing this problem exhibit very high co-infection rates. (7, 8)

At present, many diagnostic approaches offering several techniques are available for VL diagnosis such as parasitological methods including microscopic techniques, culture, immunological methods for antigen and antibody based detections and molecular techniques based on PCR methods. PCR methods have been proved to be one of the most effective approaches for VL diagnosis due to its high sensitivity and specificity, rapidity and ability to perform with a broad range of clinical specimens. The specificity of PCR can be adapted to specific need by targeting conserved regions of a gene which have high copy numbers. Moreover, the *Leishmania* species could be identified by sequencing PCR products, which could benefit for the epidemiological study of the disease. However, the obvious drawbacks of this method is mainly the need for specimens collected using an invasive methods such as, blood, bone marrow or tissue biopsies requiring equipment and specifically trained personnel to perform. (2, 9) The major advantages of using saliva rather than blood are easy access, noninvasive; expertise not needed to collect the sample and inexpensive The purpose of collecting the fluid is to detect the disease and follow up the affected individuals under treatment. (10)

In 2015, Siriyasatien et al. (11) demonstrated that the saliva would be a good source to detect *Leishmania* DNA using PCR method which could be used as a biomarker for the diagnosis of *Leishmania* infection. To determine the reliability of saliva in diagnosing *Leishmania* infection, this study aimed to detect *Leishmania* DNA in saliva specimens collected from asymptomatic *Leishmania* infection among patients with HIV/AIDS in an affected area and to evaluate the sensitivity and specificity of the nested-PCR method targeting the Internal Transcribe Space 1 (ITS1) region of the small subunit ribosomal (rRNA) gene. This study could help develop an alternative approach using a noninvasive and simpler diagnostic method for VL.

## MATERIALS AND METHODS

### Study population

A total of 305 patients with HIV/AIDS attending an HIV clinic at Trang Hospital, Trang Province were enrolled in this study between February 2015 and February 2016. Eligible participants were ≥18 years old and visited the clinic every six months for follow-up testing and received antiretroviral therapy (ART). Paired saliva and blood samples were collected from each patient to detect leishmaniasis. Demographic data of each patient were collected from face-to-face interview using a standardized questionnaire. The clinical characteristics of all patients such as clinical symptoms, underlying diseases, drug treatment as well as CD4+ levels were also collected from patients’ medical records.

### Saliva collection

Unstimulated whole saliva was collected by spitting method from each subject. Subjects were instructed to spit approximately 1-2 mL of saliva specimens into sterile collection vials. Saliva samples were stored at -20°C within 1-2 hours until used to detect *Leishmania* DNA by nested-PCR.

### Blood collection

Eight milliliters of anticoagulant (EDTA) blood specimens were collected from each patient with HIV/AIDS. The whole blood was centrifuged at 900 *g* for 10 min to separate plasma and buffy coat. Plasma and buffy coat were separately collected in micro-centrifuged tubes and kept at -20°C until used. Plasma was assayed for antibodies against *Leishmania* infection using the Direct Agglutination Test (DAT) while buffy coat was used to detect *Leishmania* DNA by nested-PCR method.

### Storage and transport

All study samples were frozen at -20°C after collecting and transported on dry ice for laboratory analysis. Samples were processed within one month at the Department of Parasitology, Phramongkutklao College of Medicine, Bangkok, Thailand.

### DNA extraction

DNA was extracted from 200 µl of each saliva sample using Gen UP™ gDNA Kit (biotechrabbit, Germany) following manufacturer instructions and with a final elution volume of 40 µl. The extracted DNA samples were kept at -20°C until used. Saliva from healthy individuals was used as negative control.

For the buffy coat samples, DNA from each sample (200 µl) was extracted using Gen UP™ gDNA Kit (biotechrabbit, Germany) following manufacturer instructions. Final elution volume of 40 µL was collected and used for DNA template.

### Leishmania DNA detection

Nested-PCR method was performed to detect the *Leishmania* DNA in extracted saliva and buffy coat samples by amplifying the ITS1 region of the rRNA gene. The primers for the primary PCR reaction were LITSR (5’-CTGGATCATTTTCCGATG-3’) and L5.8 (5’-TGATACCACTTATCGCACTT-3’) (12); while the primers for the secondary PCR reaction were LITS2R (5’-CTGGATCATTTTCCGATGATT-3’) and L5.8S inner (5’-GTTATGTGAGCCGTTATCC-3’) (13, 14).

The primary PCR reaction was conducted in a total volume of 50 µl, containing 25 pmol of each primer, 0.2 mM dNTP (Promega, USA), 1.5 mM MgCl_2_, 1x PCR buffer, 2 U of *Taq* DNA polymerase (Promega, USA) and 10 µL of DNA template (saliva)/ 4 µL of DNA template (buffy coat). The nested-PCR reaction was performed in a total volume of 25 µl, containing 25 pmol of each primer, 0.2 mM dNTP (Promega, USA), 1.5 mM MgCl_2_, 1x PCR buffer, 2 U of *Taq* DNA polymerase (Promega, USA) and 1 µL (saliva)/ 4 µL (buffy coat) of primary PCR product. The PCR fragment was amplified under the following thermocycler conditions: (i) 2 min at 95°C; (ii) 35 cycles of 20 sec at 95°C, 30 sec at 53°C and 1 min at 72°C; and (iii) 6 min at 72°C. Extracted DNA from cultured *L. martiniquensis* promastigotes (MON-229, WHO code: MHOM/MQ/92/MAR1) was used as the positive control. The size of the PCR product of the ITS1 region was 300-350 bp depending on *Leishmania* species.

PCR products from each reaction were separated in 1.5% agarose gel (Vivantis, Malaysia) containing 0.04 µg/ml SYBR^®^ Safe DNA gel stain (Invitrogen, USA). Ten µL of each PCR product was diluted with 1:5 volume of loading dye and loaded in each well of agarose gel. The 100 bp DNA ladder (Vivantis, Malaysia) was used as molecular marker to evaluate the size of amplicons. Gel electrophoresis was carried out at 100 volts for 45 min in TBE buffer. PCR product was visualized under UV light and documented by Molecular Imager^®^ Gel Doc^TM^ XR+ System with Imager Lab^TM^ 3.0 (BioRad, USA).

### Nucleotide sequence and data analysis

All positive nested-PCR products were purified and bi-directionally sequenced. The sequencing results were then validated using BioEdit Version 7. The validated sequences were compared with the reference sequences of *Leishmania* retrieved from the GenBank database to identify the species. The nested-PCR results obtained from saliva samples were compared with those obtained from buffy coat samples identically collected and processed in the same way as described.

### Detection of *Leishmania* antibodies

Detection of *Leishmania* antibodies was performed using a commercial DAT kit (DAT-KIT, Biomedical Research, Amsterdam, The Netherlands) according to manufacturer instructions. The positive plasma control was obtained from confirmed VL cases using the nested-PCR method While plasma from healthy individuals was used for the negative control. The cutoff value of positive DAT titers was ≥1:100 following manufacturer recommendation.

### Statistical analysis

The positive nested-PCR results of both saliva and buffy coat samples were compared with those of a gold standard method. We used positive *Leishmania* nucleotide sequences obtained from the nested-PCR results of saliva and/or buffy coat obtained from the subjects as the gold standard method. The sensitivity and specificity of the nested-PCR-ITS1 assays were calculated using a 2X2 contingence table on the basis of the cross distribution of negative and positive results with 95% confidence interval (CI). Characteristics of patients were set according to the results of three diagnostic methods; DAT, the nested-PCR-ITS1-saliva and the nested-PCR-ITS1-buffy coat, to determine *Leishmania* infection. The association between patients’ characteristics and *Leishmania* infection was assessed using the chi-square test and *p* values <0.05 were considered statistically significant. All analyses were performed using STATA, Version SE14 (Stata Corporation, College Station, Texas).

### Ethics statement

The research protocol was reviewed and approved by the Ethics Committee of the Royal Thai Army Medical Department and the Ethics Committee of Mahidol University, Thailand. Written informed consent was obtained from all participants included in the study.

## RESULTS

### Leishmania DNA detection

Positive *Leishmania* DNA in saliva/buffy coat samples was defined as detected DNA using the nested-PCR-ITS1 assay and positive *Leishmania* determined by nucleotide sequencing. Of 305 patients with HIV/AIDS, 176 were shown to be positive according to at least one of the nested-PCR-ITS1 methods using saliva and/or buffy coat, together with positive sequences of *Leishmania* DNA. These 176 patients were considered true positive. Positive samples were 15.1 and 42.6% by the nested-PCR-ITS1 using saliva and buffy coat, respectively. The result agreement between the nested-PCR-ITS1 using buffy coat and saliva was 52%. Antibody detection for *Leishmania* by the DAT was 43.61% (133/305) and result agreement between DAT and the nested-PCR-ITS1 using saliva was 46.5%. Agreement of positive results among the three methods; the nested ITS1-PCR using saliva, the nested ITS1-PCR using buffy coat and DAT was 12.1% as shown in Table 1.

**Table 1:** The results of *Leishmania* DNA detection by the nested-PCR-ITS1 using saliva (SL), and the nested-PCR-ITS1 using buffy coat (BF) including antibody detection using the Direct Agglutination Test (DAT), n (%)

|  | Positive | Negative |
| --- | --- | --- |
| Nested ITS1-PCR using saliva (SL) | 46 (15.1) | 259 (84.9) |
| Nested ITS1-PCR using buffy coat (BC) | 130 (42.6) | 175 (57.3) |
| DAT using plasma | 133 (43.6) | 172 (56.4) |
| **Agreement of positive results** |  |  |
| SL & BC | 14 (4.6) |  |
| SL & DAT | 8 (2.6) |  |
| SL & BC & DAT | 5 (1.6) |  |

### Nucleotide sequences and data analysis

Of 46 positive nested-PCR samples of saliva, sequence analysis reported 3 species of *Leishmania* infections, that is, 28 (60.87%) *L. siamensis*, 9 (19.57%) *L. martiniquensis*, 1 (2.17%) *L. donovani* complex and the remaining 8 (17.39 %) were unidentified species of *Leishmania*.

### Sensitivity and specificity

As shown in Table 2, the sensitivity and specificity determined by the nested-PCR-ITS1 using buffy coat were 73.86% and 100%, respectively. However, the sensitivity and specificity by the nested-PCR-ITS1 using saliva specimens were 26.14 and 100%, respectively. The sensitivity of the nested-PCR-ITS1 in buffy coat was significantly higher than those of saliva (*p*<0.01).

**Table 2:** Sensitivity and specificity by the nested-PCR-ITS1 using buffy coat (BF) and saliva (SL) specimens

| Nested ITS1-PCR |  | *Leishmania* sequence |  |  | % Sensitivity (95% CI) | % Specificity (95% CI) |
| --- | --- | --- | --- | --- | --- | --- |
|  |  | Positive | Negative | Total |  |  |
| Saliva | Positive | 46 | 0 | 46 | 26.1 | 100 |
|  | Negative | 130 | 129 | 259 | (17.7-35.7) | (96.4-100) |
| Buffy coat | Positive | 130 | 0 | 130 | 73.9 | 100 |
|  | Negative | 46 | 129 | 175 | (63.2-81.4) | (96.4-100) |

### Characteristics of participants who presented positive test results for *Leishmania* infection

The demographic data and clinical characteristics were collected from a standardized questionnaire and medical records as shown in Table 3. Of 305 patients, 285 (93.44%) questionnaires were collected. The mean age of participants was 43.2 years old, 85.6% had obtained an educational level of secondary school or lower, 12.3% were unemployed, 82.8% lived in Trang Province, 12.6% had traveled abroad, 15.1% were intravenous drug users, 9.8% had received a diagnosis of opportunistic infection, 9.2% had a CD4+ cell count less than 200 cells/mL in the past 6 months and 95.1% had undetectable viral load in the past 6 months (<50 copies). Regarding the analysis of three diagnostic methods for *Leishmania* infection; i) DNA detection by the nested ITS1-PCR using saliva; ii) DNA detection by the nested ITS1-PCR using buffy coat and iii) antibody detection using DAT, no significant difference was observed among age group, gender, educational level, province, history of going abroad and opportunistic infection. The characteristics of 31 patients who had *Leishmania* infection diagnosed by the nested-PCR-ITS1 using buffy coat and/or saliva are shown in Table 4. Variable ranges of CD4 levels (50-962 cells/μl; mean= 468.7 ± 258.3) and DAT titers (1:100-1:6400) were observed among 31 patients. Two patients exhibited symptomatic VL while the remaining patients presented asymptomatic VL. Two symptomatic VL cases showed positive results in all three assays with a high DAT titer at 1:6400. However, three asymptomatic VL patients had positive results of the three assays with the DAT titer at 1:1600, 1:400 and 1:400, respectively. Nine asymptomatic VL patients were detected for *Leishmania* DNA in both saliva and buffy coat; however, their antibodies were undetected using DAT. Only three asymptomatic VL cases exhibited positive results of both saliva and DAT methods. Additionally, 14 asymptomatic VL cases exhibited positive results of both buffy coat and DAT. Opportunistic infections were found only among four patients. Of 31 patients, 29 had undetectable viral load (<50 copies/mL). Most asymptomatic VL patients were not IDUs while opportunistic infections were found only among three patients.

**Table 3:** Characteristics of patients according to the results of three diagnostic methods for *Leishmania* infection: *i)* Antibody detection using Direct Agglutination test (DAT); *ii)* DNA detection by the nested-PCR-ITS1 using buffy coat and *iii)* DNA detection by the nested-PCR-ITS1 using saliva

| Characteristics | Total<br>n=285 | Nested ITS1-PCR using saliva |  |  | Nested ITS1-PCR using buffy coat |  |  | Antibody detection by DAT |  |  |
| --- | --- | --- | --- | --- | --- | --- | --- | --- | --- | --- |
|  |  | Negative | Positive | p-value | Negative | Positive | p-value | Negative | Positive | p-value |
| **Age (Mean and SD)** | 43.2±7.4 | 43.0±8.6 | 43.2±7.4 | 0.84 | 42.4±8.4 | 44.0±8.4 | 0.39 | 43.6±8.5 | 42.4±8.3 | 0.82 |
| **Gender** |  |  |  | 0.94 |  |  | 0.43 |  |  | 0.72 |
| Female | 145 (50.9) | 122 (50.6) | 23 (52.8) |  | 88 (53.0) | 57 (47.9) |  | 82 (50.3) | 63 (51.6) |  |
| Male | 140 (49.1) | 119 (49.4) | 21 (47.7) |  | 78 (47.0) | 62 (52.1) |  | 81 (49.7) | 59 (48.4) |  |
| **Education** |  |  |  | 0.85 |  |  | 0.89 |  |  | 0.69 |
| Primary Education | 121 (42.4) | 104 (43.2) | 17 (38.6) |  | 71 (42.8) | 50 (42.0) |  | 71 (43.6) | 50 (41.0) |  |
| Secondary Education | 123 (43.2) | 103 (42.7) | 20 (45.5) |  | 70 (42.2) | 53 (44.5) |  | 71 (43.6) | 52 (42.6) |  |
| Bachelor or higher | 41 (14.4) | 34 (14.1) | 7 (15.9) |  | 25 (15.1) | 16 (13.4) |  | 21 (12.9) | 20 (16.4) |  |
| **Occupation** |  |  |  | 0.32 |  |  | 0.49 |  |  | 0.09 |
| Planting and farming | 65 (22.8) | 58 (24.0) | 7 (15.9) |  | 33 (19.8) | 32 (26.9) |  | 44 (27.0) | 21 (17.2) |  |
| Animal handlers | 55 (19.3) | 49 (20.3) | 6 (13.6) |  | 37 (22.3) | 18 (15.1) |  | 26 (15.9) | 29 (23.8) |  |
| Business and government | 47 (16.5) | 37 (15.4) | 10 (22.7) |  | 27 (16.3) | 20 (16.8) |  | 22 (13.5) | 25 (20.5) |  |
| Laborers | 83 (29.1) | 70 (29.0) | 13 (29.5) |  | 49 (29.5) | 34 (28.6) |  | 51 (31.3) | 32 (26.2) |  |
| Unemployed | 35 (12.3) | 27 (11.2) | 8 (18.2) |  | 20 (12.0) | 15 (12.6) |  | 20 (12.3) | 15 (12.3) |  |
| **Province** |  |  |  | 0.81 |  |  | 0.27 |  |  | 0.52 |
| Trang | 236 (82.8) | 199 (82.6) | 37 (84.1) |  | 134 (80.7) | 102 (85.7) |  | 137 (84.0) | 99 (81.1) |  |
| Others | 49 (17.2) | 42 (17.4) | 7 (15.9) |  | 32 (19.3) | 17 (14.3) |  | 26 (16.0) | 23 (18.9) |  |
| **Travel abroad** |  |  |  | 0.23 |  |  | 0.46 |  |  | 0.38 |
| No | 249 (87.4) | 213 (88.4) | 36 (81.8) |  | 143 (86.1) | 106 (89.1) |  | 140 (85.9) | 109 (89.3) |  |
| Yes | 36 (12.6) | 28 (11.6) | 8 (18.2) |  | 23 (13.9) | 13 (10.9) |  | 23 (14.1) | 13 (10.7) |  |
| **Intravenous drug user** |  |  |  | 0.01 |  |  | 0.02 |  |  | 0.01 |
| No | 242 (84.9) | 210 (87.1) | 32 (72.7) |  | 148 (89.2) | 94 (79.0) |  | 131 (80.4) | 111 (91.0) |  |
| Yes | 43 (15.1) | 31 (12.9) | 12 (27.3) |  | 18 (10.8) | 25 (21.0) |  | 32 (19.6) | 11 (9.0) |  |
| **Opportunistic infection** |  |  |  | 0.47 |  |  | 0.91 |  |  | 0.11 |
| No | 257 (90.2) | 216 (89.6) | 41 (93.2) |  | 150 (90.4) | 107 (89.9) |  | 151 (92.6) | 106 (86.9) |  |
| Yes | 28 (9.8) | 25 (10.4) | 3 (6.8) |  | 16 (9.6) | 12 (10.1) |  | 12 (7.4) | 16 (13.1) |  |
| **CD4+ count (cells/mL)** |  |  |  | 0.04 |  |  | 0.65 |  |  | 0.25 |
| <200 | 26 (9.2) | 20 (8.3) | 6 (13.6) |  | 17 (10.4) | 9 (7.6) |  | 13 (8.1) | 13 (10.7) |  |
| 200-500 | 149 (52.3) | 134 (55.8) | 15 (34.9) |  | 87 (53.0) | 62 (52.1) |  | 80 (49.7) | 69 (56.6) |  |
| >500 | 108 (37.9) | 86 (35.8) | 22 (51.2) |  | 60 (36.6) | 48 (40.3) |  | 68 (42.2) | 40 (32.8) |  |
| **Viral load detection** |  |  |  | 0.39 |  |  | 0.91 |  |  | 0.01 |
| >50 copies | 13 (4.9) | 12 (5.4) | 1 (2.3) |  | 8 (5.0) | 5 (4.7) |  | 3 (2.0) | 10 (8.47) |  |
| <50 copies | 254 (95.1) | 212 (94.6) | 42 (97.7) |  | 152 (95.0) | 102 (95.3) |  | 146 (98.0) | 108 (91.5) |  |

**Table 4:** Characteristics of 31 patients, who presented positive results for *Leishmania* infection by the nested-PCR-ITS1 using saliva and/or the nested-PCR-ITS1 using buffy coat including DAT titer, CD4 level, opportunistic infection, viral load and IVDUs, are shown

| Patient # | Nested ITS1-PCR saliva | Nested ITS1-PCR buffy coat | DAT Titer | CD4+ level | Opportunistic infection | Viral load | IVDUs |
| --- | --- | --- | --- | --- | --- | --- | --- |
| 1 | + | + | Negative | 688 | No | <50 | No |
| 2 | + | + | Negative | 272 | No | <50 | Yes |
| 3 | - | + | 1:1600 | 249 | No | >50 | No |
| 4 | + | + | Negative | 424 | No | <50 | Yes |
| 5 | - | + | 1:800 | 756 | No | >50 | No |
| 6 | - | + | 1:800 | 602 | No | <50 | No |
| 7 | + | + | 1:1600 | 481 | No | <50 | No |
| 8 | + | + | Negative | 167 | No | <50 | No |
| 9 | + | + | 1:400 | 561 | No | <50 | No |
| 10 | + | - | 1:800 | 172 | No | <50 | No |
| 11 | + | + | Negative | 594 | No | <50 | Yes |
| 12 | + | + | Negative | 838 | No | <50 | Yes |
| 13 | - | + | 1:800 | 560 | No | <50 | No |
| 14 | + | + | Negative | 668 | No | <50 | No |
| 15 | + | - | 1:800 | 50 | No | >50 | No |
| 16 | + | + | Negative | 843 | TB | <50 | Yes |
| 17 | - | + | 1:800 | 838 | No | <50 | No |
| 18 | - | + | 1:1600 | 216 | No | <50 | No |
| *19 | + | + | 1:6400 | 259 | No | <50 | No |
| 20 | + | + | Negative | 42 | No | <50 | No |
| 21 | - | + | 1:100 | 965 | TB, PUL | <50 | No |
| 22 | - | + | 1:400 | 352 | No | <50 | No |
| 23 | - | + | 1:400 | 878 | TB, PUL | <50 | No |
| 24 | - | + | 1:400 | 196 | TB, PUL | <50 | No |
| 25 | - | + | 1:400 | 376 | No | <50 | No |
| 26 | + | - | 1:800 | 286 | No | <50 | No |
| 27 | + | + | 1:400 | 246 | No | <50 | Yes |
| 28 | - | + | 1:400 | 519 | No | <50 | No |
| 29 | - | + | 1:100 | 463 | No | <50 | No |
| 30 | - | + | 1:200 | 346 | No | <50 | No |
| *31 | + | + | 1:6400 | 622 | No | <50 | Yes |

Note; TB= Tuberculosis, PUL= Pulmonary infection, \*Symptomatic VL

## DISCUSSION

Saliva is increasingly being considered as a potential noninvasive alternative to blood for *Leishmania* diagnosis. Sample collection and simple handling methods that do not disrupt the quality of samples are ideal in field settings in developing countries. This study was conducted to determine the diagnosis performance of nested-PCR using saliva, compared with nested-PCR using buffy coat to detect *Leishmania* infection. The ITS1 region of the SSU-rRNA gene was chosen as a target for diagnostic PCR assay for *Leishmania* DNA together with the nested-PCR to increase the sensitivity of the test (15-17). As a result, no concordance was revealed for the majority of positive results obtained among the salivary, buffy coat as well as plasma antibodies. Due to low immunity status among patients positive for HIV, many factors should be taken into account, i.e., CD4+ levels, viral load, nutritional status, hormonal status and opportunistic infection including severity of *Leishmania* infection. Levels of parasites circulating in the blood might be involved in time-related secretion of *Leishmania* DNA in saliva. Unfortunately, in this study, quantification of *Leishmania* parasites in the buffy coat among these asymptomatic VL patients with HIV was not investigated. Thus, a relationship between *Leishmania* DNA in saliva and severity of the disease using level of parasitemia could not be demonstrated. Our results showed that *Leishmania* DNA was detected in saliva before being found in blood circulation in a number of asymptomatic VL patients with HIV and vice versa; a number of buffy coat samples showed positive nested-PCR but not in saliva. This could be due to a very low number of *Leishmania* parasites in the blood circulation during early infection when parasites have not yet infiltrated the salivary glands.(18) The rK39 strip test (InBios, Seattle, WA) was used to detect the presence of antibodies against rK39 to confirm VL infection in sputum samples.(18) The confirmed VL patients from bone marrow smears showed a relationship between antibodies in serum and sputum samples. The antibodies are carried from blood to saliva through the diffusion process (19-21). Our study also showed positive result agreements for DAT, nested-PCR of buffy coat and saliva in two symptomatic or clinical VL patients. Thus, saliva could be a good choice of clinical specimens to diagnosed merely symptomatic VL and for treatment follow-up of the disease. Secretion of pathogens from blood to saliva and time-related secretions of infectious agents might differ among organisms, i.e., viruses, bacteria, fungi and protozoa. When saliva was tested among patients with HIV, saliva was superior to both serum and urine regarding sensitivity and specificity of symptomatic patients (10). Studies on HIV infection showed the advantage when saliva can be used as a reliable diagnostic specimen. Using a saliva collection commercial kit to detect *Plasmodium falciparum* DNA in saliva, high sensitivities of 95% and 100% were observed between nested-PCR-saliva and nested-PCR-blood, respectively.(22) In contrast to enzyme secretion in saliva regarding malaria infection a study showed a weak correlation between levels of lactate dehydrogenase of *P. falciparum* in saliva and blood parasitemia (23). Little information is known on the factors associated with *Leishmania* in blood and saliva; thus, more studies of these clinical samples during disease progression of VL are required.

In this study, the sensitivity of the nested PCR-ITS1 using unstimulated whole saliva was 26.1%, which was much lower than the nested PCR-ITS1 using buffy coat (73.9%). This low sensitivity of nested-PCR in unstimulated whole saliva samples without any preservatives could be a result of the effects of sample storage temperature and a time delay of 1 to 2 hours before keeping saliva samples in -20°C. From the point of collection, DNA stability could be affected during storage condition especially in saliva samples containing small amounts of parasite DNA. Thus, buffy coat remains a good clinical specimen to screen *Leishmania* DNA in the blood of both symptomatic and asymptomatic VL among immunocompetent and immunocompromised individuals. To increase the sensitivity to detect *Leishmania* DNA, two clinical specimens, buffy coat and saliva, should be simultaneously used for laboratory diagnosis.

The patients participating in this study were individuals with HIV/AIDS. The results of this study showed that detectable viral load (>50 copies /mL) among patients with HIV/AIDS could also elicit *Leishmania* antibody production revealing significantly differences of positive DAT between those with a detectable viral load and nondetectable viral load. Related studies have shown that positive antibody by DAT did not show concordance with positive *Leishmania* DNA in the blood among patients with HIV/AIDS (14).

Among 31 patients with HIV/AIDS presenting *Leishmania* infection, 29 asymptomatic and 2 symptomatic VL cases had agreement with positive results using either two or three assays (Table 4). Two symptomatic VL cases showed concordance of positive nested-PCR-saliva, nested-PCR-buffy coat as well as DAT titer at 1:6400. However, early detection of *Leishmania* DNA is very beneficial to the remaining 29 asymptomatic VL cases because clinical symptoms have not yet presented. So far, treatment protocols for asymptomatic VL cases have not been recommended. Thus, a follow-up study of these patients using both saliva and buffy coat should be performed to determine the progression of infection to clinical VL or remission from the infection caused by *Leishmania* parasites. The results of this study also agreed with the published paper by Chusri et al., 2012, which studied symptomatic HIV/VL among Thai patients. They found positive *Leishmania* DNA in both buffy coat and saliva and follow-up cases were performed before and after VL treatment of which *Leishmania* DNA was eliminated and disappeared after the treatment (24). As a result, they noted the usefulness of saliva to diagnose specimens in both symptomatic and asymptomatic HIV/VL cases. Due to a small number of cases in a related study, the sensitivity and specificity could not be calculated. The results of this study revealed a lower sensitivity of the nested-PCR-ITS1 using saliva when compared with that of the nested-PCR-ITS1 using buffy coat.

This study also confirmed the presence of at least three species of *Leishmania* infections, that is, *L. siamensis*, *L. martiniquensis* and *L. donovani* complex. Of these, *L. siamensis* was the most predominant species, followed by *L. martiniquensis* which were identified among asymptomatic VL patients with HIV/AIDS in Trang Province, southern Thailand. Moreover, public health awareness for disease transmission should be of extreme concern because infected persons could spread the disease in other areas.

In conclusion, this study demonstrated that the nested-PCR-ITS1 assays using saliva samples should not be routinely used to early detect *Leishmania* infection especially, among patients with asymptomatic VL due to its lower sensitivity compared with those using buffy coat. At present, buffy coat appears to be a better specimen by far to diagnose asymptomatic and symptomatic VL infection among individuals with HIV/AIDS. The use of both buffy coat and saliva together as clinical specimens would increase the sensitivity of *Leishmania* detection. Further improvement for saliva collection i.e., samples frozen from the point of collection before DNA extraction or the use of a saliva collection kit to preserve the parasite DNA could offer an opportunity to use saliva to diagnose VL.

## ACKNOWLEDGEMENTS

We would like to express our sincere gratitude to all enrolled participants who provided clinical samples and information for the questionnaire. We also wish to thank the staff of the HIV Clinic, Trang Hospital, Trang Province for their fruitful cooperation. This research was supported by the National Science and Technology Development Agency, Science and Achievement Scholarship of Thailand and Phramongkutklao College of Medicine Research Fund.

## REFERENCES

1. WHO. 2017. Leishmaniasis Fact Sheet, *on* WHO. http://www.who.int/gho/neglected_diseases/leishmaniasis/en/. Accessed 10th October.

2. Singh S. 2006. New developments in diagnosis of leishmaniasis. Indian J Med Res 123:311–30.

3. Kevric I, Cappel MA, Keeling JH. 2015. New World and Old World Leishmania Infections: A Practical Review. Dermatol Clin 33:579–93.

4. Herwaldt BL. 1999. Leishmaniasis. Lancet 354:1191–9.

5. Leelayoova S, Siripattanapipong S, Manomat J, Piyaraj P, Tan-Ariya P, Bualert L, Mungthin M. 2017. Leishmaniasis in Thailand: A Review of Causative Agents and Situations. Am J Trop Med Hyg 96:534–542.

6. van Griensven J, Ritmeijer K, Lynen L, Diro E. 2014. Visceral leishmaniasis as an AIDS defining condition: towards consistency across WHO guidelines. PLoS Negl Trop Dis 8:e2916.

7. Lindoso JA, Cunha MA, Queiroz IT, Moreira CH. 2016. Leishmaniasis-HIV coinfection: current challenges. HIV AIDS (Auckl) 8:147–156.

8. Alvar J, Aparicio P, Aseffa A, Den Boer M, Canavate C, Dedet JP, Gradoni L, Ter Horst R, Lopez-Velez R, Moreno J. 2008. The relationship between leishmaniasis and AIDS: the second 10 years. Clin Microbiol Rev 21:334–59, table of contents.

9. Blackwell JM. 1992. Leishmaniasis epidemiology: all down to the DNA. Parasitology 104 Suppl:S19–34.

10. Streckfus C, Bigler L. 2002. Saliva as a diagnostic fluid. Oral diseases 8:69–76.

11. Siriyasatien P, Chusri S, Kraivichian K, Jariyapan N, Hortiwakul T, Silpapojakul K, Pym AM, Phumee A. 2016. Early detection of novel Leishmania species DNA in the saliva of two HIV-infected patients. BMC Infect Dis 16:89.

12. El Tai NO, El Fari M, Mauricio I, Miles MA, Oskam L, El Safi SH, Presber WH, Schonian G. 2001. Leishmania donovani: intraspecific polymorphisms of Sudanese isolates revealed by PCR-based analyses and DNA sequencing. Exp Parasitol 97:35–44.

13. Untergasser A, Cutcutache I, Koressaar T, Ye J, Faircloth BC, Remm M, Rozen SG. 2012. Primer3—new capabilities and interfaces. Nucleic acids research 40:e115–e115.

14. Manomat J, Leelayoova S, Bualert L, Tan-Ariya P, Siripattanapipong S, Mungthin M, Naaglor T, Piyaraj P. 2017. Prevalence and risk factors associated with Leishmania infection in Trang Province, southern Thailand. PLoS Negl Trop Dis 11:e0006095.

15. Schönian G, Nasereddin A, Dinse N, Schweynoch C, Schallig HD, Presber W, Jaffe CL. 2003. PCR diagnosis and characterization of Leishmania in local and imported clinical samples. Diagnostic microbiology and infectious disease 47:349–358.

16. Bensoussan E, Nasereddin A, Jonas F, Schnur LF, Jaffe CL. 2006. Comparison of PCR assays for diagnosis of cutaneous leishmaniasis. J Clin Microbiol 44:1435–9.

17. Reuss SM, Dunbar MD, Calderwood Mays MB, Owen JL, Mallicote MF, Archer LL, Wellehan JF, Jr. 2012. Autochthonous Leishmania siamensis in horse, Florida, USA. Emerg Infect Dis 18:1545–7.

18. Singh D, Pandey K, Das VN, Das S, Kumar S, Topno RK, Das P. 2009. Novel noninvasive method for diagnosis of visceral leishmaniasis by rK39 testing of sputum samples. J Clin Microbiol 47:2684–5.

19. Sharma U, Singh S. 2009. Immunobiology of leishmaniasis. Indian J Exp Biol 47:412–23.

20. Singh S. 2014. Changing trends in the epidemiology, clinical presentation, and diagnosis of Leishmania–HIV co-infection in India. International Journal of Infectious Diseases 29:103–112.

21. Pfaffe T, Cooper-White J, Beyerlein P, Kostner K, Punyadeera C. 2011. Diagnostic potential of saliva: current state and future applications. Clinical chemistry 57:675–687.

22. Mfuh KO, Tassi Yunga S, Esemu LF, Bekindaka ON, Yonga J, Djontu JC, Mbakop CD, Taylor DW, Nerurkar VR, Leke RGF. 2017. Detection of Plasmodium falciparum DNA in saliva samples stored at room temperature: potential for a non-invasive saliva-based diagnostic test for malaria. Malar J 16:434.

23. Nambati EA, Kiarie WC, Kimani F, Kimotho JH, Otinga MS, Too E, Kaniaru S, Limson J, Bulimo W. 2018. Unclear association between levels of Plasmodium falciparum lactate dehydrogenase (PfLDH) in saliva of malaria patients and blood parasitaemia: diagnostic implications? Malar J 17:9.

24. Chusri S, Hortiwakul T, Silpapojakul K, Siriyasatien P. 2012. Consecutive cutaneous and visceral leishmaniasis manifestations involving a novel Leishmania species in two HIV patients in Thailand. Am J Trop Med Hyg 87:76–80.

